# Quantifying the Emergence of Population-Level Activity in Neuronal Systems

**DOI:** 10.64898/2026.02.13.705719

**Authors:** Hardik Rajpal, Pedro A.M. Mediano, Madalina I. Sas, Henrik J. Jensen, Fernando E. Rosas

**Affiliations:** Centre for Complexity Science, Imperial College London, London, United Kingdom; Department of Mathematics, Imperial College London, London, United Kingdom; I-X Centre for AI in Science, Imperial College London, London, United Kingdom; Department of Computing, Imperial College London, London, United Kingdom; Department of Psychology, University of Cambridge, Cambridge, United Kingdom; Department of Brain Sciences, Imperial College London, London, United Kingdom; Department of Informatics, University of Sussex, Brighton, United Kingdom; Centre for Eudaimonia and human flourishing, University of Oxford, Oxford, United Kingdom

## Abstract

Collective neural phenomena, such as oscillations and avalanches, are high-level neural signatures observed in aggregated spiking neuronal activity which have been consistently associated with a range of cognitive functions. However, it is often hard to elucidate whether such phenomena are mere epiphenomena or have causal or informational relevance. In this work, we investigate this question by leveraging recent information-theoretic tools to identify emergent phenomena between relevant scales of neural activity. For this, we propose a computational framework combining information-theoretic and network science principles, which we use to investigate emergence in both in-vivo datasets and in-silico simulations. Our approach enables characterisation of emergence phenomena, identifies the relevant scales at which they take place, and elucidates the network-level mechanisms that underpin them. Results show that in-vivo neuronal oscillations show substantial emergent behaviour for smaller prediction delays, while avalanches maintain their emergent nature for larger timescales. These results are supported by in-silico simulations, which show that the emergent signature of oscillations is facilitated by the network structure and interneuronal time-delays. Overall, these results highlight the role of network-level interactions between groups of neuronal assemblies as the key driver of emergent population activity in the brain.

## 1 Introduction

Collective neural phenomena, such as oscillations and avalanches, are thought to be crucial for various cognitive functions^1,2^ such as sensory processing^3^, memory consolidation^4,5^, and decision making^6,7^. However, whether these population-level activities are causally-relevant emergent phenomena or just epiphenomena of the underlying neural activity is still a matter of debate^8^. On one hand, various theoretical accounts have argued for the causal role of oscillations and other aggregated neural activity in the function and self-organization of the brain^9–14^. On the other hand, other studies have argued that any causal role attributed to population-level activity is confounded by various factors, such as the geometry^15^, shared distributed inputs^16^, inconsistent spike-field relationships^17,18^, and overall a lack of identifiable biological causal mechanisms. This tension between the two views has been discussed in detail in recent reviews^19,20^.

Alongside theoretical developments, there has been a growing body of empirical work investigating the emergence of population-level activity in the brain and its relevance for behaviour and cognition. Experimental studies have established correlations between spectral properties of oscillations and various aspects of different cognitive tasks^21,22^. More recently, causal interventions using optogenetics^23^, transcranial alternating current (tACS)^24^, electrical (tECS)^25^, and magnetic (tMS)^26^ stimulation have provided evidence supporting the causal role of oscillations in cognitive functions. Similarly, the presence of neuronal avalanches has been associated with optimal information processing in the brain^27–30^, and interventions that disrupt critical dynamics have been shown to impair cognitive functions^31–33^.

Computational models have complemented these empirical findings, providing additional insights into the role of population-level activity in the brain. Various models of spiking neuronal networks have been shown to exhibit oscillations and avalanches under different conditions^34–40^. These models have advanced our understanding of the mechanisms that drive the emergence of these phenomena, such as the role of network structure, synaptic plasticity, and neuronal dynamics. Furthermore, they have been used to develop quantitative measures to identify statistical causal measures of emergence^41–43^ and efficient neural network architectures^44^.

Despite these advances, several challenges remain in understanding and quantifying emergence in the brain. Firstly, in order to quantify emergence from neural data, we need to identify the relevant scales of neuronal activity that give rise to population-level phenomena. Neuronal activity is often highly redundant, with groups of neurons exhibiting similar firing patterns due to shared inputs and local connectivity^45–47^. This redundancy can obscure the identification of emergent phenomena, and can result in spurious identification of emergence^48^. Therefore, it is crucial to identify the relevant scales of neuronal activity that capture the underlying dynamics while minimizing redundancy. Once the relevant scales are identified, we need robust measures that can quantify the relationships between these scales. Such quantitative measures can be used to address key questions related to the emergence of population-level activity in neuroscience. First, does the population-level activity possess enhanced predictability as compared to the micro-level activity of the neurons? Second, does the population-level activity exert a downward influence by conditioning the future activity of the neurons? Lastly, what are the mechanisms that drive the emergence of population-level activity?

Recently, information-theoretic measures of emergence have been developed to directly quantify emergence and tackle the above questions^49–51^. These measures provide a way of quantifying the excess information in a supervinient macro feature of the system as compared to the information stored in the parts of the system (in the Granger-causal sense^52^). While these metrics can be computed efficiently for systems with a relatively small number of sub-components, they struggle to provide reliable estimates for large systems due to the ‘curse of dimensionality’. To address this limitation, practical measures that can be estimated using marginal and pairwise mutual information have been proposed^50^. These practical measures quantify the difference between the information provided by the macro feature about a target as compared to the parts of the system. Despite their advantages, these practical measures tend to lose statistical power to detect emergence in large systems, as they double-count redundant information among the parts. To address this issue, improved estimators of emergence have been recently proposed that correct for higher-order redundancies^53^. Correcting for the shared redundancy is crucial for biological systems, where redundancy is a useful feature that provides robustness^54,55^.

Here, we leverage these improved estimators of causal emergence and network-based dimensionality reduction to develop a computational framework to investigate emergence in neuronal systems. We use this framework to analyse both a dataset of neuronal recordings from mouse visual cortex^56^ and also a computational model of Izhikevich spiking neurons connected in a Pyramidial Inter-Neuronal Gamma (PING) architecture. Our results show how this approach allow us to (a) identify the relevant scales of neuronal activity (assemblies) that give rise to population-level phenomena, (b) quantify the presence and nature of emergence in population-level activity, and (c) identify the network-level mechanisms that drive the emergence using computational models.

## 2 Results

### 2.1 Detecting emergence in neural activity

We sought to investigate the conditions under which oscillations are emergent and exert a downward causal power on the activity of neurons. For this, we analyzed an existing dataset of neuronal recordings from mouse visual cortex^56^ that displays oscillations during sleep and avalanches during wakefulness. We complement this analysis by also studying a computational model of Izhikevich spiking neurons^35^.

We deploy a two-step redundancy minimizing pipeline to identify the relevant scales of neuronal activity in both experimental and simulated spike trains. Our approach follows three steps:

(i) We use a network-based community detection algorithm to identify neuronal assemblies from the pairwise mutual information network of the spike trains. The identified communities represent the neuronal assemblies which are highly correlated and carry shared redundant information.
(ii) We then use the community-level spike counts as the primary (micro) scale of our analysis, which reduces the dimensionality of the data and enables a more accurate estimation of emergence measures.
(iii) We finally quantify information-theoretic emergence in both experimentally recorded and simulated spiking activity using the improved estimators of emergence measures that account for higher-order redundancies in the system.

The measures used in the last step are the three practical measures of *causal emergence* (Ψ), *downward causation* (Δ) and *causal decoupling* (Γ) introduced in Ref.^50^. In brief, Ψ quantifies the amount of excess information in the macro feature about its future as compared to the information provided individually by the parts of the system. A positive value of Ψ indicates that the macro-feature is more than the sum of its parts, hence causally emergent. Similarly, Δ quantifies the maximum excess information provided by the macro feature about the future activity of any of the parts as compared to information provided by the parts. Positive values of Δ indicate a greater statistical influence of the macro feature in constraining the future activity of the parts. Finally, Γ = 0 is sufficient to conclude that the macro feature provides no information about the future of any of the parts and is causally decoupled from the dynamics of the parts. That said, please note that Δ *>* 0 or Γ = 0 only lead to the above conditions when Ψ *>* 0 also holds. Technical details about these measures can be found in *Methods*.

Thus, our approach is to compare the population-level neuronal spiking activity with the activity of neuronal assemblies (see Figure 1). Focusing on neuronal assemblies allows us to group neurons that are highly correlated, which enables a more accurate estimation of emergence measures via improved estimators^53^. The neuronal assemblies are identified empirically using community detection on the mutual-information network of the neuronal spike trains. Then, the population-level activity is simply the sum of the activity of the underlying neural assemblies (see Figures 1a and 1b for a schematic representation). Further details about the dataset and the model are provided in the *Methods* section.

**Figure 1.**
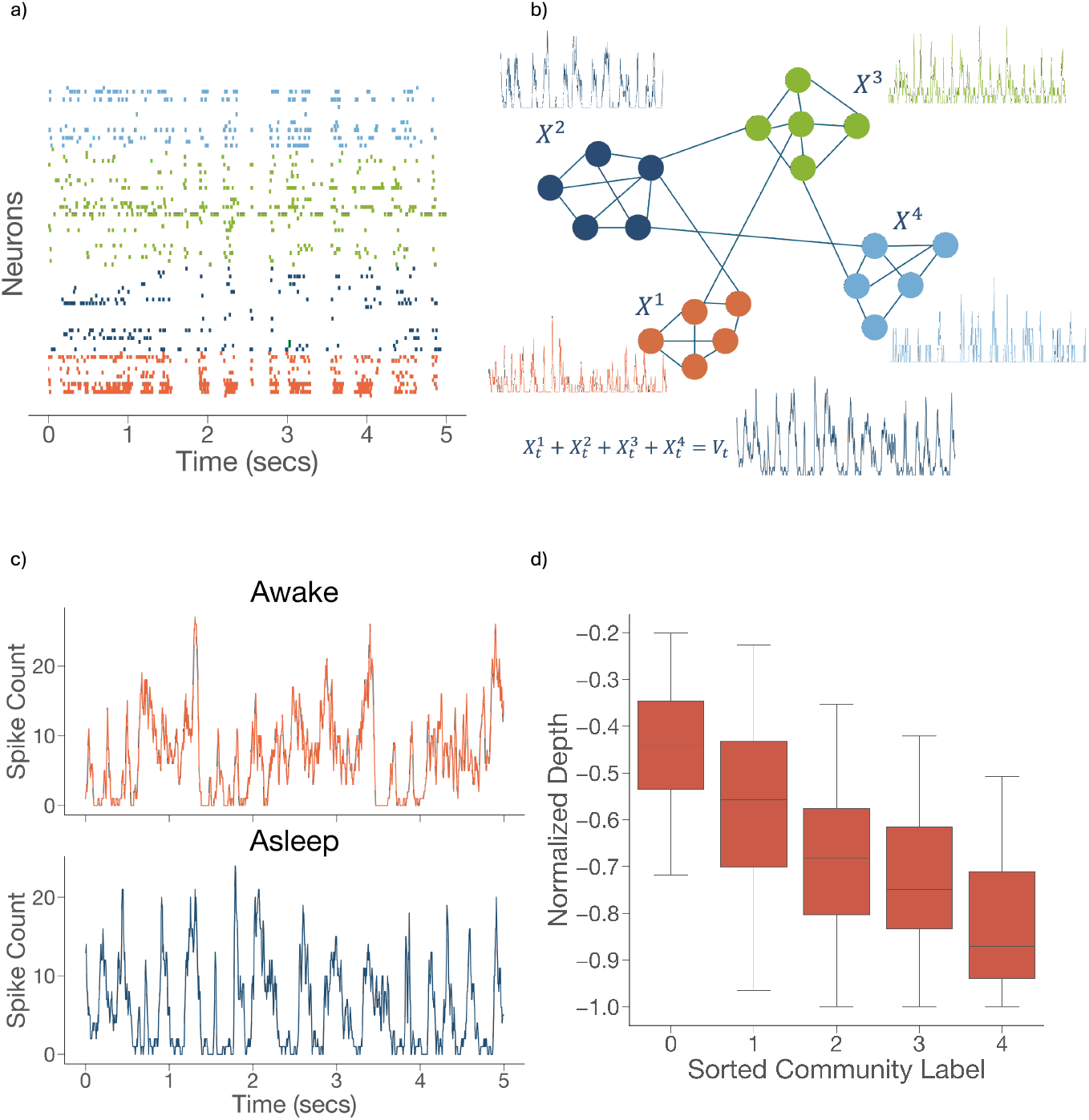
(a) Schematic representation of extracting neuronal assemblies from spiking data. (b) Pairwise mutual information is estimated using spike trains of all pairs of neurons to build a symmetric network. After extracting the backbone network, Louvain community detection is deployed to identify communities representing neuronal assemblies. Community-level spike counts (*X*_*i*_(*t*)) and population-level activity (*V* (*t*)) can then be extracted to study the relationship between the two scales and quantify emergence. (c) population-level spiking activity observed in the mouse visual cortex in two states. The top panel shows the activity in the awake state, where each period of increased spiking activity can be considered as a neuronal avalanche. It can be seen that avalanches of all possible sizes can occur in the awake state (see ^57^ for a detailed analysis). The bottom panel shows oscillations in spiking activity in the asleep state, and a strong periodic behaviour is observed with a characteristic time period. (d) Communities of neurons exhibit characteristic depth in the cortex. Normalized depths of each neuron are plotted against the community labels (sorted by depth) across the 13 experiments. The depth of each neuron is normalized by the maximum recorded depth in that experiment.

### 2.2 Emergence in the mouse visual cortex

We used the above pipeline to study emergence on a dataset consisting of recordings across 13 experiments involving mice in both awake and asleep states, consisting of neuronal spike trains recorded from a silicon probe with 64 channels spanning all cortical layers^56^. We extracted community- and population-level spiking activity from the spike-trains recorded from the mouse visual cortex during awake and asleep states^56^. The resulting neuronal assemblies are found to be depth-specific, as communities of neurons tend to be localized to a particular layer in the cortex (see Figure 1 (d)). This finding is in line with the layer-specific population activity discussed in the paper describing the datasets^56^.

We consider the resulting community-level spike counts as the activity observed at the micro scale of the system, and regarded the total population level activity as macro-level activity. The population level activity during the sleep state can be seen to exhibit oscillatory behaviour, while periods of low and high activity appear intermittently in the awake state (see Figure 1 (c)). The population-level activity observed in the awake state has been shown to be associated with critical avalanches. No characteristic duration or size of the avalanches is observed in the population-level activity, which is a signature of self-organized criticality in the awake state^57^.

Using the time series of spiking activity of the parts and the whole population, we compute the three practical measures of emergence defined in a previous publication^50^. We use the improved estimators of emergence measures that account for higher-order redundancies in the system^53^ and correct for the estimation bias by comparing to surrogate data generated by randomly shifting the spike-trains. This bias-correction enables us to account for the spurious positive values that may occur in non-interacting systems^48^. Leveraging these corrections, we estimate the three measures for different prediction delays *τ* between the community-level and the macro activity.

Our results reveal that both causal emergence and downward causation take place at both awake and sleep conditions. The average values of surrogate corrected measures over the 13 experiments are shown in Figure 2. Interestingly, the metrics are higher for oscillatory population activity than for avalanches; it is significantly larger than zero for longer prediction delays for the latter. In particular, emergence only takes place in the range of 0.2 − 0.3 seconds for the awake condition, which relates to the dominant frequency of 3 − 6 Hz, which is known to be important during the awake state^56^. These enhanced positive values of the emergence measures in this range were not observed in the asleep state.

**Figure 2.**
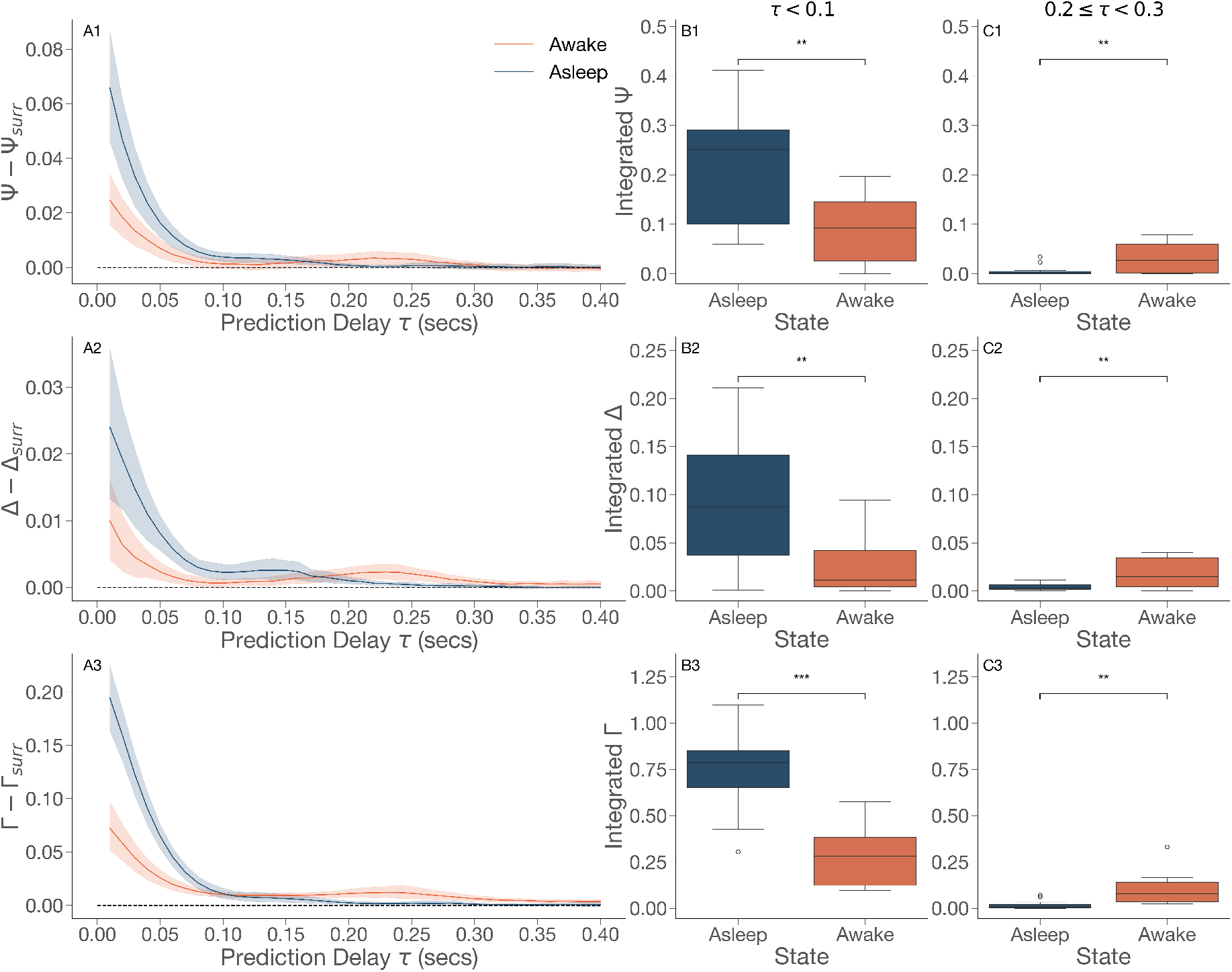
(A)The three emergence measures are plotted after surrogate correction for the different prediction time delays for the Awake and Asleep states. (B) For shorter timescales (*τ <* 0.1), the integrated area under the curve of causal emergence Ψ and downward causation Δ are significantly higher in the asleep (oscillatory) state. (C) However, for longer timescales (0.2 ≤ *τ <* 0.3), the integrated area under the curve of Ψ and Δ is higher in the awake state. (A3) Γ is found to be non-zero throughout the relevant range of prediction delays when other causal measures are positive, indicating the absence of causal decoupling.

Additionally, our results did not show evidence for causal decoupling, as Γ was found to be significantly above zero only at ranges at which the system showed causal emergence.

### 2.3 Emergence in the Izhikevich model

To identify the conditions under which neural oscillations can be emergent and to identify the network properties that modulate them, we study a model of Izhikevich neurons connected in a Pyramidial Inter-Neuronal Gamma (PING) architecture. Groups of excitatory and inhibitory neurons connected in a PING architecture have shown synchronized oscillatory behaviour for spontaneous input^36^. In order to draw parallels with the dynamics of neuronal assemblies discussed in this paper, we discuss a variant of PING architecture which has two communities of excitatory neurons representing the neuronal assemblies at the superficial and deeper layers of the visual cortex, respectively. The excitatory neurons in each community are connected to a common inhibitory population, which regulates the excitatory activity.

The network structure of the excitatory neurons is generated using a stochastic block model (SBM) with two communities representing the superficial and the deep layers of the cortex. It can be observed that the connectivity structure of the inter-layer interactions modulates the oscillatory behaviour of the population activity (see Figure 3). Therefore, the connectivity parameters of the SBM are chosen to reflect the functional connectivity of neurons observed in the mouse visual cortex during awake and asleep states. In previous work, it has been shown that during deep sleep, the feedforward connections from the superficial layer to the deeper layers weaken while the recurrent connections within the deeper layers are strengthened. To model this, we simultaneously decrease the connection probability from the superficial layer to the deeper layer (*p*_*cross*_) and increase the connection probability within the deeper layer (*p*_*deep*_), while keeping all other parameters constant. Since inter-neuronal time delays significantly impact the frequency of oscillatory behaviour in PING architectures, we sample the time-delays from a uniform distribution between 1 and *D*_*max*_. Neuronal spiking activity is simulated on these networks using the standard parameters of Izhikevich neurons for different values of the connection probabilities and time delay distribution between the excitatory neurons. The network is simulated for 5 seconds, and the recorded spiking activity is used to compute the Ψ and Δ measures of emergence discussed above. The obtained emergence measures are corrected for the estimation bias using the surrogate datasets generated using a null network where all excitatory connections were set to zero.

**Figure 3.**
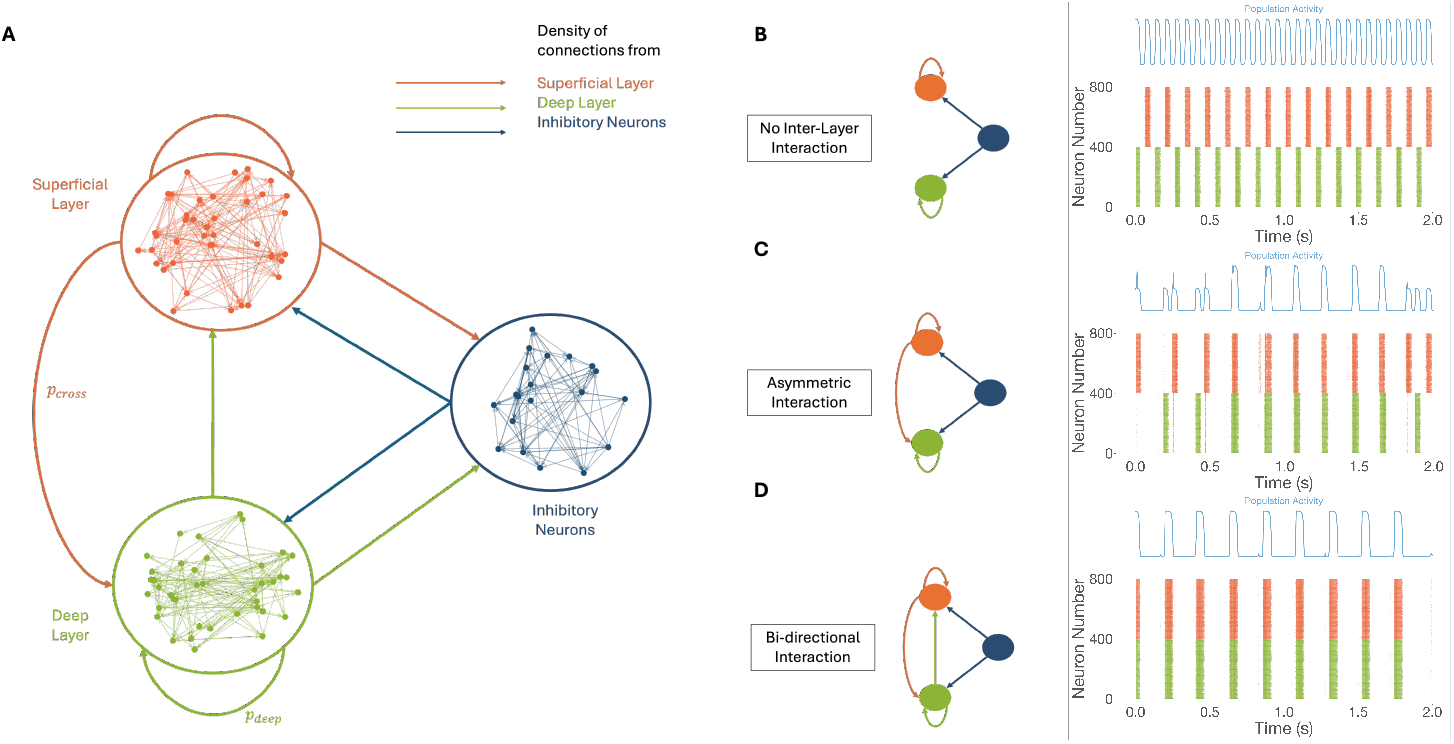
**A** Schematic representation of the computational model of Izhikevich spiking neurons connected in a PING architecture. The model consists of two communities of excitatory neurons representing the neuronal assemblies at the superficial and deeper layers of the visual cortex, respectively. The excitatory neurons in each community are connected to a common inhibitory population, which regulates the excitatory activity. By varying the inter-layer connectivity structure, the model can simulate activity with a varying degree of synchronization among the layers. **B** It can be seen that when the layers are disconnected, they exhibit an alternating oscillatory activity, and the population activity has half the time period of the individual layers. **C** In case of asymmetric connectivity, we observe a metastable state where layers sometimes spike together but can get desynchronized over time. **D** When the layers are connected bi-directionally, they exhibit synchronized oscillatory activity with a larger characteristic time period. The population activity has the same time period as the individual layers.

First, we investigate the impact of inter-neuronal time delays on the emergence measures. Our results show that both Ψ and Δ measures are positive for larger values of *D*_*max*_, which characterizes the range of time delays in the network (see Figure 4). For smaller values of *D*_*max*_, although Ψ is positive, it decreases to a minimum and then increases again for larger values of *D*_*max*_. On the other hand, Δ is found to be increasing for larger values of *D*_*max*_. This indicates that a wide range of heterogeneous time delays supports both emergence and downward causation of synchronized neuronal oscillations.

**Figure 4.**
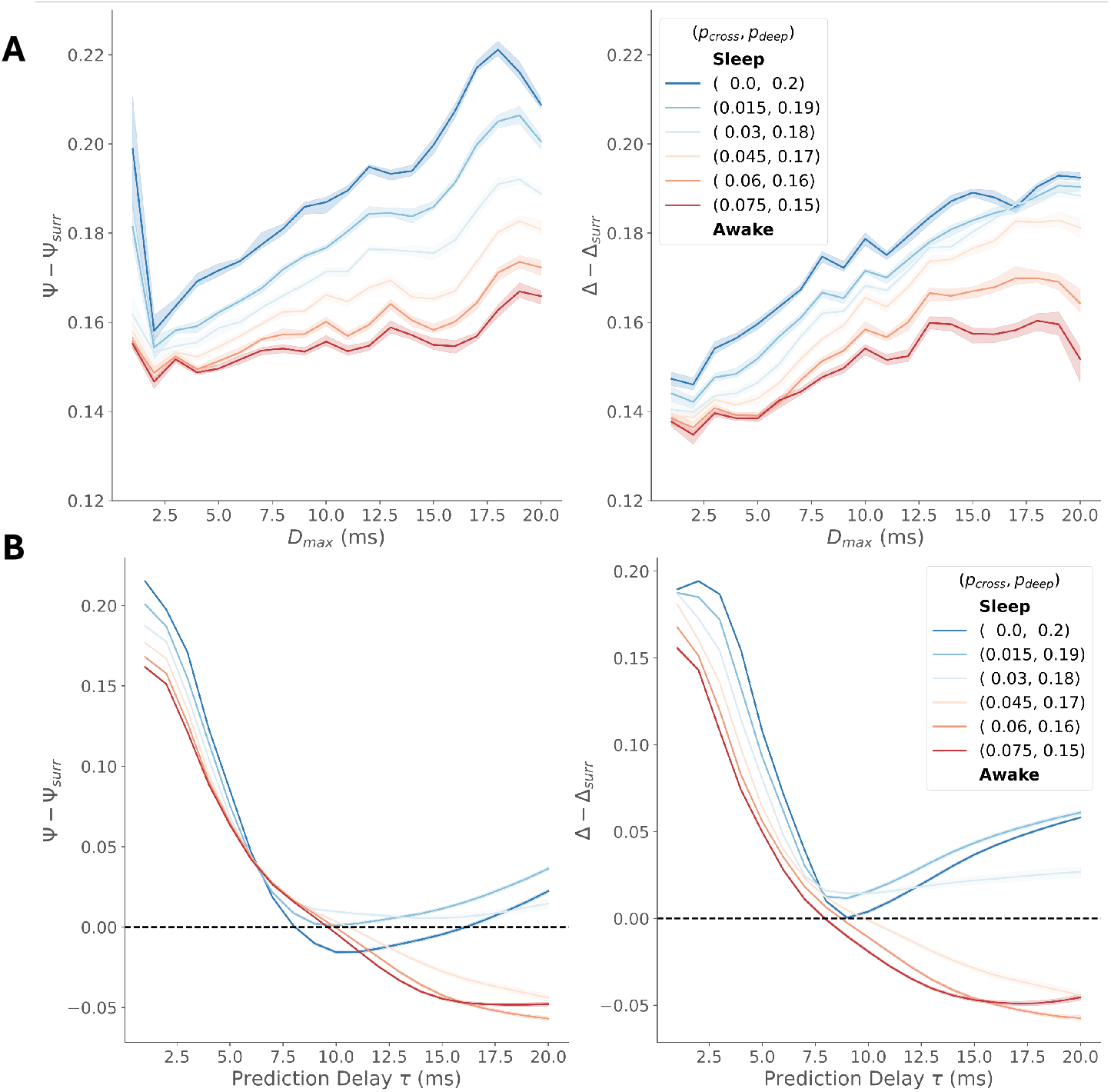
**(A)** Values of the emergence measures Ψ and Δ for different values of time delay distributions charactrized by *D*_*max*_. The measures are averaged over 200 different networks for each value of *D*_*max*_. **(B)** Values of the emergence measures for different values of connection probabilities *p*_*cross*_ and *p*_*deep*_ from superficial to the deeper layers. It must be noted that as *p*_*cross*_ is varied from 0.075 to zero, simultaneously *p*_*deep*_ is increased from 0.15 to 0.2. The measures are averaged over 100 different networks for each combination of *p*_*cross*_ and *p*_*deep*_ with *D*_*max*_ ∈ (15, 21].

Focusing on large ranges of time delays (*D*_*max*_ *>* 15 ms), we investigate the impact of awake-asleep neuronal connectivity on the emergence measures. By simultaneously decreasing *p*_*cross*_ and increasing *p*_*deep*_, we can observe the impact of deep sleep on the emergence of oscillations. It can be seen that both Ψ and Δ are greater for smaller values of *p*_*cross*_ for a short prediction horizon *tau*. When comparing *p*_*cross*_ = 0 and *p*_*cross*_ = 0.015, we see that as emergence measures decrease for low values of tau (for increased *p*_*cross*_), they comparatively increase for larger prediction horizons. These results mimic the results observed in the experimental data, where the emergence measures were found to be positive for larger time delays in the awake state. However, beyond this value of *p*_*cross*_, we found that measures decrease for all values of *tau* when *p*_*cross*_ is increased. Thus, the model supports the experimental findings that the emergence horizon of the population activity is dependent on the connectivity between the excitatory neurons in the two layers.

## 3 Discussion

In this work, we have developed a computational framework to quantitatively investigate emergent phenomena in spiking neural activity, and we have applied this framework to both in-vivo data and a computational model. Our results highlight the role of network interactions at the scale of groups of neuronal assemblies as the key driver of the emergence of population activity in the brain. The analysis of the experimental data crucially highlighted the differences in the strength and range of emergence measures in the awake and asleep states. The awake state exhibited a greater range of prediction delays for which the population-level activity was emergent, while the asleep state exhibited a strong oscillatory behaviour with a characteristic time period. The computational model of Izhikevich spiking neurons connected in a PING architecture supported these findings, as we varied the network structure to recreate the interactions observed in the experiments among neuronal assemblies. The model also highlighted the role of heterogeneous inter-neuronal time delays and symmetry in network structure in modulating the emergence of oscillations.

The findings suggest that different network structures and interaction mechanisms can drive the system into dynamical regimes where the population activity exhibits collective emergent behaviour. Under these conditions, the different dynamical modes of activity are distributed across neuronal assemblies, which coordinate together to give rise to the emergent population activity. This higher-order coordination not only facilitates enhanced predictability of the population activity but also exerts a downward causal influence on the future activity of the neuronal assemblies. These findings provide insights into the network-level mechanisms that drive the emergence of population-level activity in the brain, and highlight the importance of considering higher-order interactions and network structure in understanding brain function.

The computational framework presented here provides tools to quantify emergence in biological systems where there exists local redundancy for resilience in the system, while having global coordination across the different parts of the system. The combination of network-based dimensionality reduction and the improved estimators of emergence measures provides a powerful tool to quantify the relationship between different scales of neuronal activity. The framework can be applied to other large systems where emergence is a key feature alongside widespread redundancy, such as in social networks, ecological systems and other biological complex systems.

The results presented here using the model are focused on understanding the interactions that drive neuronal oscillations. However, the framework can be extended to study other emergent phenomena in neuronal activity, such as avalanches and criticality. This would involve simulating neuronal networks with state-time dependent plasticity or other known mechanisms that have shown self -organized criticality in the brain^58,59^. The framework can also be applied to other types of neuronal models, such as integrate-and-fire models or conductance-based models, which provide a way to simulate other biologically relevant population activity patterns. Having shown the applicability of the framework to the case of oscillations, we can now extend it to other non-trivial cases of emergence where these measures can be used as an exploratory tool to identify population activity patterns that correspond to specific cognitive states.

## 4 Methods

The results presented in the paper are based on information-theoretic analysis of neuronal spiking activity from experimental recordings and computational models. The experimental data is obtained from the mouse visual cortex during awake and asleep states^56^. The computational model is based on Izhikevich spiking neurons connected in a PING architecture with two communities of excitatory neurons. The neuronal assemblies are identified using community detection on the mutual information network of the spike trains to identify the relevant scales of neuronal activity in both systems. The emergence measures are estimated, between the scales of global activity and the identified neuronal assemblies, using the improved estimators that account for higher-order redundancies in the system^53^.

### 4.1 Experimental data

The experimental data is obtained from the mouse visual cortex during awake and asleep states^56^. The dataset includes neuronal spike trains recorded from a silicon probe (64-channel, linear, single shank, Cambridge NeuroTech H3 64×1 probe) spanning all cortical layers. Each experiment involves recordings from mice in both awake and asleep states. The dataset provides the location of each neuron (67±18 units per recording) in the cortical layers along with the raw spiking activity associated with each neuron. The raw data is processed to extract the neuronal spike trains at the millisecond resolution. The dataset also provides information about the state of the subjects during the course of multi-hour recordings (6.2±1.8), which are classified as awake, non-REM sleep and REM sleep. For the analysis presented in this paper, we focus on the awake and non-REM sleep states. We analyzed 13 of the 19 available datasets, as the remaining datasets did not have at least 30 minutes of continuous recordings in either of the two states. The dataset is publicly available online (https://buzsakilab.com/wp/database/), and for further details please refer to the original publication^56^.

### 4.2 Computational model

The computational model is based on Izhikevich spiking neurons connected in a PING architecture with two communities of excitatory neurons modulated by a population of inhibitory neurons. The model is implemented in Python using the standard parameters of the Izhikevich model^35^. The neurons are connected in a Pyramidial Inter-Neuronal Gamma (PING) architecture with two communities of excitatory neurons representing the neuronal assemblies at the superficial and deeper layers of the visual cortex, respectively. The excitatory neurons in each community are connected to a common inhibitory population, which regulates the excitatory activity (see Figure 3). The network structure of the excitatory neurons is generated using a stochastic block model (SBM) with two communities.

The parameters of the SBM are chosen to reflect the functional connectivity of neurons observed in the mouse visual cortex during awake and asleep states. In previous work, it has been shown that during deep sleep, the feedforward connections from the superficial layer to the deeper layers weaken while the recurrent connections within the deeper layers are strengthened. To model this effect, we simultaneously vary the connection probability from the superficial layer to the deeper layer (*p*_*cross*_) and the connection probability within the deeper layer (*p*_*deep*_), while keeping all other parameters constant. The inter-neuronal time delays are sampled from a uniform distribution between 1 and *D*_*max*_, where *D*_*max*_ is varied to study the impact of heterogeneous time delays on the emergence of oscillations.

Once the network is generated, the spiking dynamics are simulated using the differential equations described in the Izhikevich model^35^.

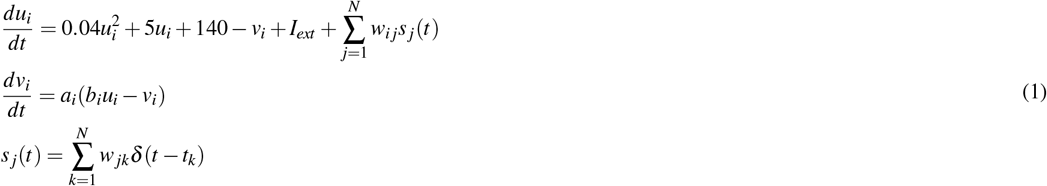

where *u*_*i*_ is the membrane potential of neuron *i, v*_*i*_ is the recovery variable, *I*_*ext*_ is the external input to the neuron, *w*_*i j*_ is the synaptic weight from neuron *j* to neuron *i*, and *s* _*j*_(*t*) is the spike train of neuron *j*. The parameters *a*_*i*_, *b*_*i*_ are specific to each neuron and are chosen to reflect the dynamics of excitatory and inhibitory neurons in the PING architecture. When the membrane potential *u*_*i*_ reaches a threshold, the neuron spikes and the recovery variable *v*_*i*_ is reset. The time delay between the spikes of the neurons is incorporated in the model by using a delay line for each synapse, which introduces a delay in the transmission of spikes from one neuron to another.

The spiking activity on a given network is simulated by numerically integrating the differential equations using the Euler method with a time step of 1 ms. The simulation is run for 5 seconds, and the spiking activity is recorded at each time step. The recorded spiking activity is then used to compute the emergence measures discussed above.

### 4.3 Quantifying emergence

The emergence measures are computed using the recorded spiking activity from both the experimental data and the computational model. The neuronal assemblies are identified using community detection on the mutual information network of the spike trains to identify the relevant scales of neuronal activity in both systems. The community-level spike counts are computed as the sum of spikes from all neurons in a community at each time step. The population-level activity is defined as the sum of community-level spike counts across all communities at each time step. We then quantify information-theoretic interactions between the community-level activity and the population-level activity using the measures of causal emergence (Ψ) and downward causation (Δ).

Mutual Information (MI) is used to quantify the statistical dependence between any to time series *X* (*t*) and *Y* (*t*), defined as:

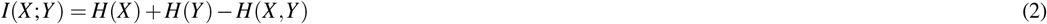

where *H*(*X*) and *H*(*Y*) are the entropies of the time series *X* (*t*) and *Y* (*t*) respectively, and *H*(*X,Y*) is the joint entropy of the two time series. In the context of this paper, *X* (*t*) is the community-level spike counts and *Y* (*t*) is the population-level activity. We discretize the spike counts into logarithmically spaced bins to compute the entropies and mutual information. The Shannon entropy of a discrete random variable *X* is defined in terms of its probability distribution *p*(*x*) as:

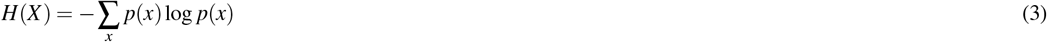

First, we estimate the pairwise mutual information between all pairs of neuronal spike trains to estimate the dependence network of the neurons. Then we deploy the Louvain community detection algorithm^60^ on the mutual information network to identify the communities of neurons representing the neuronal assemblies. The community-level spike counts are computed as the sum of spikes from all neurons in a community at each time step 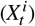. The population-level activity is defined as the sum of community-level spike counts across all communities at each time step 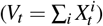. The community-level spike counts and the population-level activity are then used to compute the emergence measures.

The measures of emergence are defined in terms of mutual information by Rosas et al. (2020)^50^ as follows,

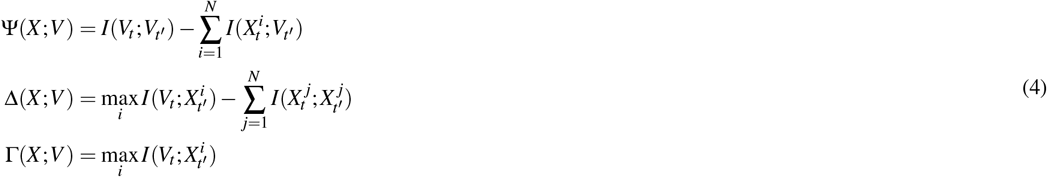

The measure Ψ(*X;V*) quantifies the excess information in the population level activity *V* about its future as compared to the information provided by the community level activity *X*^*i*^. A positive value of Ψ indicates that the population level activity is more than the sum of its parts, hence emergent. Similarly, Δ(*X;V*) quantifies the maximum excess information provided by the population level activity about the future activity of any of the communities as compared to information provided by the communities. Positive values of Δ indicate a greater statistical influence of the population level activity in constraining the future activity of the communities. Finally, Γ(*X;V*) = 0 indicates that the population level activity provides no information about the future of any of the communities and is causally decoupled from the dynamics of the communities. The emergence measures are estimated for different prediction delays *τ* between the community-level and the population-level activity (*t*^*′*^ = *t* + *τ*). The emergence measures are corrected for the estimation bias using the surrogate datasets generated by randomly shifting the spike-trains of each neuron, which breaks any temporal correlations in the data. The emergence measures are then estimated on these surrogate datasets to obtain a baseline for comparison. It has been shown that a positive Ψ is observed in a system of non-interacting random walkers^48^. Therefore, it is important to compare the values obtained from these measures to an appropriate null distribution.

#### 4.3.1 Shared Redundancy Corrections

However, the emergence measures as defined above tend to underestimate the amount of emergence in the system, as they double-count the shared redundant information among the parts 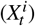 when estimating Ψ(*X;V*) and Δ(*X;V*). To correct for this bias, we use the improved estimators of emergence measures that account for higher-order redundancies in the system^53^, which iteratively add the shared redundancy terms to the emergence measures. In our analysis, we consider the first-order (pairwise) and second-order (triplet) redundancies in the system. Theemergence measures with the first-order corrections for Ψ and Δ can be written as follows,

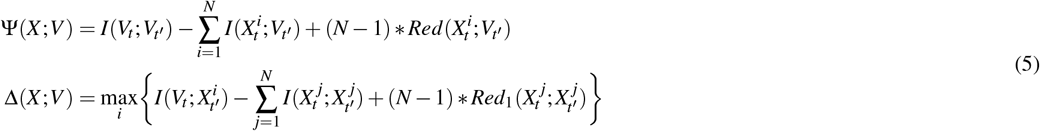

where 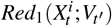 is the first-order redundant information among the community level activity 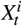 and the population level activity 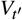. The amount of redundant information can be estimated by using an appropriate redundancy function (CITE). In our work, we use the minimum mutual information as the redundancy function, which is defined as:

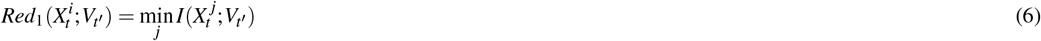

As discussed in previous work^53^, the first-order corrections provide the maximum contribution to the corrections of the emergence measures. However, in some cases, the second-order corrections can also be significant, especially when the system exhibits higher-order redundancies. We use the second-order corrections in estimating the emergence measures when there are more than two communities identified in the system. The second-order corrections can be written as follows,

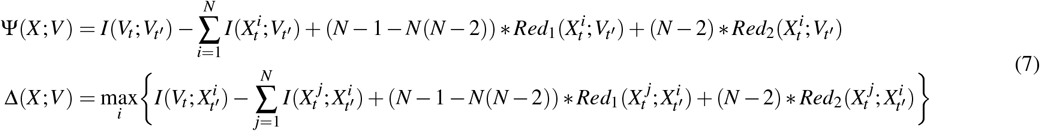

where 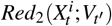is the second-order redundant information among the community level activity 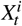 and the population level activity 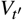. The second-order redundant information can be estimated by using the first-order redundant information among the pairs of communities, which is defined as,

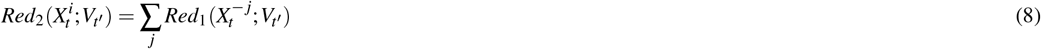

where 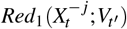 is the first order redundancy among the population level activity and the subset of community level activity excluding the *j*^*th*^ community. The total second-order correction 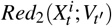 is estimated iteratively by summing the first-order redundancies by excluding one community at a time. In this study, we restrict ourselves to second-order corrections. For a detailed derivation of *k*-th order corrections, please refer to the original publication^53^. We used the Java Information Dynamics Toolkit (JIDT)^61^ to estimate the mutual information and the emergence measures. A Python implementation to estimate the emergence measures with corrections is available on GitHub.

